# Inconsistencies of empirical ecological network inference are governed by considerations of statistical approaches and dimensions of input data

**DOI:** 10.1101/2023.07.13.548816

**Authors:** Erik Kusch, Malyon D. Bimler, F. Guillaume Blanchet, James A. Lutz, Alejandro Ordonez

## Abstract

Identifying the most suitable method of ecological network inference in line with individual research considerations is a non-trivial task, which significantly hinders adoption of network approaches to forest management applications. To advance the study of ecological networks and better guide their use in managing forest ecosystems, we propose a framework that aligns pairwise species-association inference methods with specific research questions, biological interaction types, data availability, and spatial scales of study. We motivate the adoption of this framework through an empirical comparison of multiple inference methods, highlighting substantial inconsistencies that arise across scales and methodologies. Using data on species distributions and attributes at local, regional, and continental scales for temperate conifer forests in North America, we show that network inference varies significantly depending on whether occurrence, abundance, or performance data are used and the degree to which confounding factors are accounted for. Across four widely used and/or cutting-edge inference methods (COOCCUR, NETASSOC, HMSC, NDD-RIM), we find notable disparities in both whole-network metrics and pairwise species associations, particularly at continental scales. These findings underscore that no single method is likely to universally outperforms others across scales, emphasizing the importance of choosing an inference approach that aligns with specific ecological and spatial contexts. Our framework aids in interpreting network topologies and interactions in light of these method- and datatype-driven variances, providing a structured approach to more reliably infer ecological associations and address complex network dynamics in forest management practices.

## 1. Introduction

Climate change, habitat loss, and biological invasions are reshaping Earth’s biosphere (Blowes et al. 2019), including global forest communities (Allen et al. 2010), where altered species interactions (Kunstler et al. 2012) challenge ecosystem functioning (Woodward et al. 2010), forest resilience (Verbesselt et al. 2016; Liu et al. 2019) and contemporary management practices (Tang and Gustafson 1997; Allen et al. 2010; Rist and Moen 2013). Consequently, ecological and forestry research increasingly emphasizes the need to consider species interactions when predicting the emergence of novel communities and changes in ecosystem behaviour and functioning (Gilman et al. 2010). Elemental to this paradigm shift are ecological networks, which encode species identities as nodes and pairwise species interactions as links (Morales-Castilla et al. 2015) and predictions of how these nodes and links would be altered following changes in composition of ecosystems and biological communities (Kusch and Ordonez 2023).

Traditionally, studies of ecological networks have focused on local-scale interactions (Lavorel and Garnier 2002), but recent efforts have expanded to regional (Hackett et al. 2019) and even global meta-networks (Fricke et al. 2022), revealing broader-scale interaction patterns critical for forest ecosystems. In response to this trend, various inference approaches have been developed to predict interspecies interactions from species-specific input data, even where direct interaction data from field measurements or observations are unavailable (e.g., Griffith et al. 2016, Blonder and Morueta-Holme 2017, Tikhonov et al. 2020, Momal et al. 2020). However, these inference approaches encapsulate different ecological interpretations of interactions - such as mutualistic, competitive, or trophic links between species. Whereas trophic networks typically infer biological interactions as directed links (i.e., individuals of different species are influencing each other), plant-plant networks usually focus on biological association expressed as undirected links (i.e., two species occur at the same location but it is not clear whether they are interacting). While many inference approaches have been suggested for undirected link inference (e.g., Griffith et al. 2016, Blonder and Morueta-Holme 2017, Tikhonov et al. 2021), only few have been proposed for the inference of directed links (Freilich et al. 2018; Bimler et al. 2023). Consequently, forest researchers and managers need to be aware of inference approach specific link-types when selecting ecological network inference approaches for their purposes.

In addition to the distinction between direct and undirect links, methods of network inference can also be categorised along two additional data-driven axes: (1) the type of biodiversity data that is used to represent species characteristics and (2) the capacity to integrate additional data sources such as environmental conditions (Figure 1). First, biodiversity inputs and attributes which inform the inference of species interactions/associations may encode relevant information either as species occurrence, abundance, or performance – forming a spectrum of information content ranging from occurrence to performance. This spectrum establishes three distinct categorisations. Here, network inference methodologies focus on (1) occurrence (Veech 2013) or performance either implicitly through (2) abundance (Ulrich and Gotelli 2010) or (3) explicitly through biomass or fitness measurements (Bimler et al. 2023). This occurrence-performance axis arguably represents one of the most important philosophical and methodological discriminating characteristics between network inference approaches. Second, the capacity to account for different covariates such as environmental gradients, phylogenetic relatedness, and functional traits of species carries great implications for network inference performance (Morales-Castilla et al. 2015; Kusch and Vinton 2023). Approaches along this spectrum can be sorted based on covariates, like environmental conditions, being either absent or present (again, either implicitly or explicitly) when inferring the interactions/associations in a network. Most ecological network inference methodology does not incorporate such information (Blanchet et al. 2020), most likely due to data availability constraints, concerns of collinearity, or complexity of analyses. Despite the proliferation of network inference approaches for various purposes, there thus exists grounds for classification and investigation of these statistical methods to gain a better understanding of their applicability for the study and management of forest environments which has, thus far, remained unexplored. While ultimately network inference approaches should be selected based on their accuracy, comparing the accuracy of different methods requires extensive parameterisation of complex and computationally expensive simulation-validation efforts (Kusch and Vinton 2023), precluding universally applicable choices of network inference approaches at this point in time. In lieu of such wide-ranging quantitative comparisons of inference approach accuracy, a framework to support judgement of inference approach suitability and applicability for specific study purposes is prudent, timely, and within reach.

**Figure 1.**
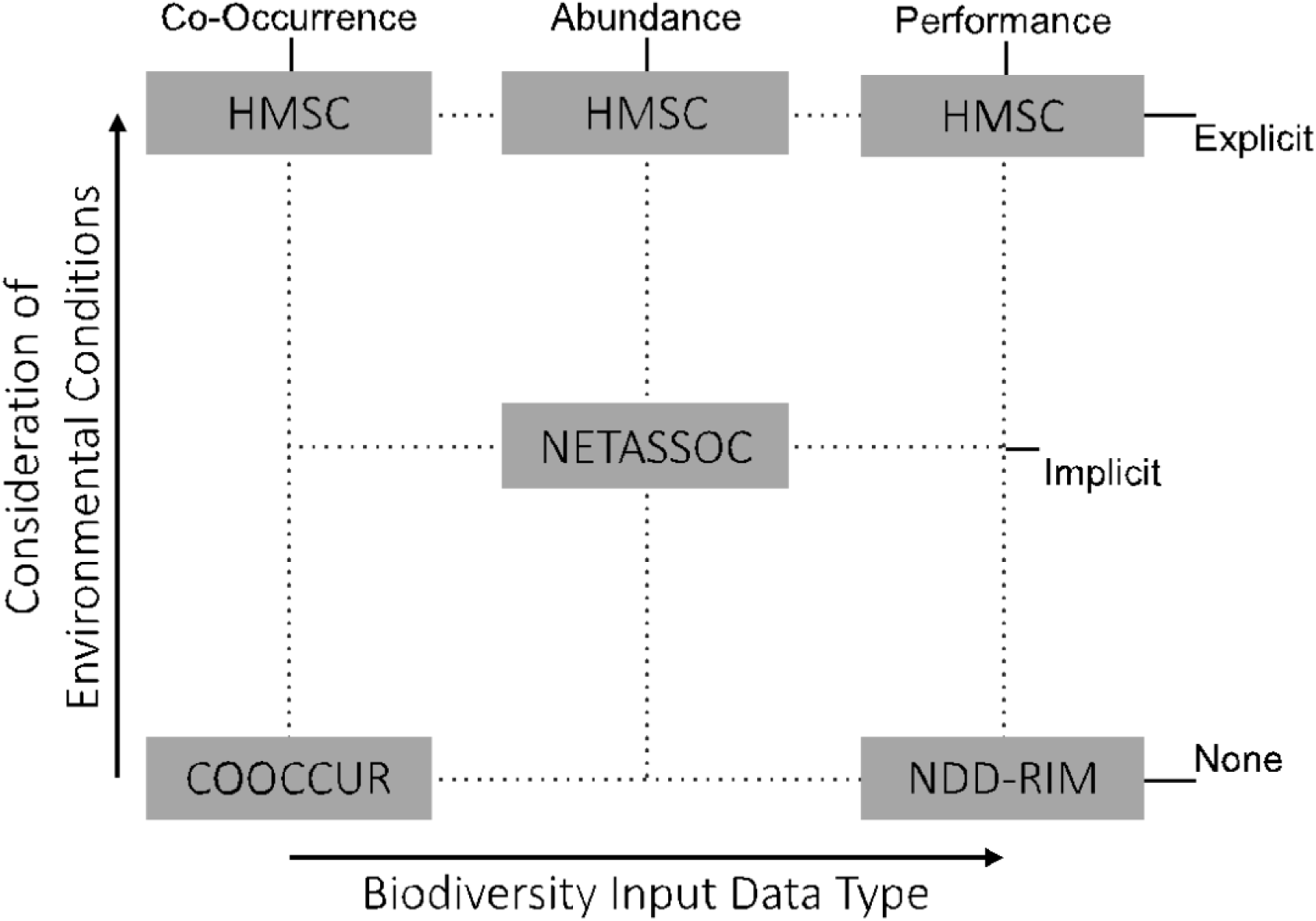
Performance-Environment Ordination of Ecological Network Inference Approaches. Network inference methodology can be distinguished with regards to their biodiversity input data type (corresponding to the occurrence-performance spectrum, see Figure S1) and their consideration of environmental conditions. The network inference approaches we selected for our study were chosen due to their comprehensive coverage of the resulting spectrum of data type use.

Delineating such a framework for choice of network inference approach with respect to link-type and data axes classifications would prove useful for the global community of forest ecosystem researchers and managers facing the decision of how to infer ecological networks for their diverse study systems at varying scales. Firstly, forest ecosystems contain both biological interactions as well as biological associations (Bello et al. 2017; Damgaard and Weiner 2021), requiring network inference approaches capable of rendering these. Secondly, forest ecosystems are understood and managed vastly differently when being assessed for species richness (e.g., management for biodiversity), biodiversity as weighted by abundance of individual species, or through measures of wood production or biomass (e.g., management for forestry and production; Rist and Moen 2013). Thirdly, the impacts of including additional covariates in analyses of ecological networks has been demonstrated by climatological conditions altering the persistence of competing species (Usinowicz and Levine 2021) and the role of functional traits in governing interactions between forest tree species (Kunstler et al. 2012). Consequently, this framework will help answering the pressing question: *How comparable are different methods of ecological network inference in real-world applications*? The necessary comparisons of modern ecological network inference approaches at scales relevant to forest management remain absent so far.

Here we establish such a framework for inference method choice in forest research and management to facilitate deeper understanding of how to best implement ecological network approaches to forestry. To do so we compare and quantify similarities between four methods of ecological network inference when inferring pairwise-associations among woody forest species across local, regional, and continental scales. Our analysis focuses on two main areas:

1. assessing consistency in network metrics across inference methods at the same scale and
2. evaluating consistency across spatial scales using the same method. We hypothesize that networks inferred using occurrence versus performance data will show structural differences, with greater similarity among methods using similar input data types. Furthermore, we anticipate that differences between approaches that account for environmental gradients and those that do not will increase at broader scales, where these gradients are primary drivers of species distribution (Stephan et al. 2021). Finally, we expect network structures to vary across scales, with spatial changes in species pools contributing to network dissimilarity but unable to fully explain the observed discrepancies (Carstensen et al. 2014). Synthesising the outcomes of this empirical consistency analysis, we conclude with a framework to guide choice of network inference approach for the study of plant-plant components of forest systems across scales.

## 2. Material & Methods

### 2.1. Ecological Network Inference Methodology

We inferred horizontal undirected (i.e., non-trophic) ecological plant-plant networks using four methods (COOCCUR, NETASSOC, HMSC, and NDD-RIM) that encapsulate a considerable variability in statistical principles, data requirements, and ecological interpretations/linktypes (Table 1). A verbose description of each inference method is provided in the supplementary material. These methods have been used primarily to infer ecological associations between species-pairs at scales ranging from local to continental applications. While other inference methods have been established (e.g.: Damgaard et al. 2018, Swain et al. 2021, Lamonica et al. 2021), we have selected this set of methods as distinct placements across the performance-environment ordination we propose for classification of such approaches (Figure 1).

**TABLE 1.** Methods of Network Inference. Description of statistical principles, data requirements, and ecological interpretation of outputs of the four methods of network inference used in this study. The evaluated methods are: **Probabilistic Species Co-Occurrence Analysis in R** (COOCCUR), **Inference of Species Associations from Co-Occurrence Data** (NETASSOC), **Hierarchical Model of Species Communities** (HMSC), and **Neighbour-Density Dependent - Response Impact Model** (NDD-RIM). In our implementation of NETASSOC, range estimates for local-scale data were not used as they did not represent informative null models. In our implementation of HMSC, trait means for each species were only used locally as functional trait expressions become too diverse at regional scales to be expressed with a mean value.

| Method | Principle | Input | Output | Interpretation of Links |
| --- | --- | --- | --- | --- |
| **COOCCUR**<br>(Griffith et al. 2016) | Probability of species-pair co-occurrence being higher or lower than random. | Site-by-Species occurrence | Standardised effect sizes (differences between expected and observed frequency of co-occurrence) | Associations, Spatial aggregation/disaggregation of species-pairs |
| **NETASSOC**<br>(Blonder and Morueta-Holme 2017) | Frequency of species-pair co-occurrence compared to null model. | Site-by-Species occurrence or abundance<br>Site-by-Species expectation of range estimates as occurrence | Standard partial correlations between species-occurrence pairings rescaled by mean and standard deviation of random generations or site-by-species expectations |  |
| **HMSC**<br>(Tikhonov et al. 2021) | Hierarchical modelling of community assembly processes were residual species-species correlations are treated as association metrics. | Site-by-Species occurrence, abundance, or performance<br>Environmental conditions at each site<br>Phylogeny of species<br>Trait means for each species | Residual correlations between species-pairing in joint distribution model | Associations, Aggregation/disaggregation (in space, abundances, or performance) of species-pairs that aren't explained through other community assembly processes |
| **NDD-RIM**<br>(Bimler et al. 2023) | Neighbour-density dependent regression and deconstruction of species interactions into response and effect coefficients for inference of rare interactions. | Individual/species performance at each site<br>Site-by-Species matrix of abundance | Regression model coefficients describing how a change in abundance of one species affects the performance of another species | Interactions, Change in performance of interaction subject with increase in abundance of interaction actor |

Data handling, network inferences, and analyses were carried out in R (R Core Team 2021). The packages cooccur (Griffith et al. 2016), netassoc (Blonder and Morueta-Holme 2017), and hmsc (Tikhonov et al. 2021) were used for the COOCCUR, NETASSOC, and HMSC methods, respectively. NDD-RIM was implemented via custom R code (see data availability statement). The workflow diagrams of data handling and network inference are presented in Figures S2 and S3.

### 2.2. Data Sources & Data Handling

We inferred networks based on two datasets of North American woody plants: the Forest Inventory Analysis Database (FIA, Burrill et al. 2018) and the Yosemite Forest Dynamics Plot (YFDP, Lutz et al. 2012). These were aggregated to describe species at three scales: local (0.256 km^2^), regional (3,030.17 km^2^), and macro (1,508,451 km^2^) representing existing scale- consideration in forest management of stand and landscape-scales (Tang and Gustafson 1997).

The local scale pool of species was defined using the YFDP data, which is a 25.6 ha study area within which all trees ≥1 cm diameter at breast height (dbh) are measured according to the protocols of the Smithsonian ForestGEO network (Davies et al. 2021). We used a subset of the YFDP data set containing 34,589 records of dbh of 11 woody plant species measured in a grid of 640, 20 m × 20 m quadrats which we subsequently treated as individual plots. For consistency with the FIA, we considered only those stems ≥2.5 cm dbh (Gray et al. 2012), retaining the records of 34,444 individuals and their dbh, as a measurement of resource allocation to growth, for network inference. These data were further aggregated to express species’ abundances and occurrence in the 640 quadrants of the YFDP.

Species pools at the regional and macro scales were defined by aggregating FIA-plot data. This dataset summarizes information on tree species’ presence, abundance, and size/biomass across the forested portion of the United States of America (Gray et al. 2012). To remove possible biases in inferred association matrices introduced by rare species, we removed all species present in less than 10% of all locations for our analyses. We used the FIA-contained records of biomass per species per plot as proxies for species’ performances. The regional scale species pool was defined by limiting FIA-plot data to the region encompassing Yosemite National Park as defined by the U.S. National Park Service (National Park Service 2009). This selection resulted in 291 observations (i.e., biomass and abundance of a species at an FIA-plot) at 101 plots containing 13 species. The macro scale species pool was defined by limiting FIA-plot data to the Temperate Conifer Forests Biome (World Wildlife Fund 2020). This masking resulted in 96,169 observations across 46,328 plots containing 15 species.

#### 2.2.1. Functional Trait Data

The YFDP dataset also includes species-specific measurements of multiple functional traits. We selected specific leaf area, leaf carbon content, and leaf nitrogen content for our analyses. These traits describe important trade-offs in plant investment in photosynthetic organs and storage of resources (Wright et al. 2004). The FIA dataset does not record functional trait measurements for its respective scales. While functional trait data is available for our target species from functional trait repositories like BIEN (Enquist et al. 2009), functional leaf trait expression is modulated by the environment (Wright et al. 2005), potentially leading to an inaccurate representation of local realities at large scales (Violle et al. 2012). We therefor chose to use functional trait means exclusively when implementing HMSC at local scale (Table 1).

#### 2.2.2. Environmental Data

To quantify plot-level environmental conditions experienced by species/individuals, we matched air temperature, soil moisture in a depth of 0-7 cm, total precipitation, and potential evaporation to the spatial scale of our data using the KrigR package (Kusch and Davy 2022). We chose these climate parameters due to their link to vegetation performance (Kusch et al. 2022). To reduce biases introduced by climate regime changes, we aggregated environmental conditions to 10-year climatologies leading up to the sampling date of each plot.

#### 2.2.3. Phylogenetic Relatedness

We established phylogenetic relatedness by pruning a phylogenetic tree supplied with the V.PhyloMaker package (Jin and Qian 2019). Species not represented in this backbone phylogeny were removed from our analyses which was in part why at the local, regional, and macroecological scale the above-mentioned 11, 13, and 15 species were respectively selected.

#### 2.2.4. Species Distributions

We obtained species range maps (for use with NETASSOC, Table 1) using the BIEN package (Maitner et al. 2018) to inform null expectations of species co-occurrences at FIA scales (i.e., regional and macro). At the YFDP scale, null models for NETASSOC were established by resampling the observed sites-by-species matrix.

### 2.3. Network Consistency Quantification

To quantify dissimilarities of inferred ecological networks at the level of individual links (interactions/associations) and node sets (species composition), we calculated association beta-diversity (a relative measure of how many present species-species links and/or species identities are shared between two networks) using the betalink (Poisot et al. 2012) R package. This comparison evaluates whether link patterns or species turnover drive changes in the identity and sign of links. We also implemented a comparison of inferred network matrices accounting links that are absent and present links thus calculating an additional inference dissimilarity score which takes into account absent associations between species shared across scales (which the betalink implementation of network beta-diversity does not). For this analysis, we reduced link sets to statistically significant links (p-value < 0.05 for COOCCUR and NETASSOC; 90% credible interval not overlapping 0 for HMSC and NDD-RIM) in the inferred weighted ecological networks. To ensure comparability across methods, we collapsed opposing links in NDD-RIM networks into mean links when computing topological metrics and assigned statistical significance only when both underlying directed links were identified as statistically significant. Additionally, we changed link weights to -1 and 1 based on their original sign to mimic large-scale decision-making information for forest inventory management of whether species-pairs suggest facilitation or competitive exclusion. The pairwise associations common to all inferred networks owing to the shared species are highlighted with black outlines in Figures 2 and 3. Subsequently, we were able to assess network inference consistency of only significantly inferred non-absent links as well as entire link-sets including non-statistically significant (thus absent) inferred links across diverging species pools and scales of assessment.

**Figure 2.**
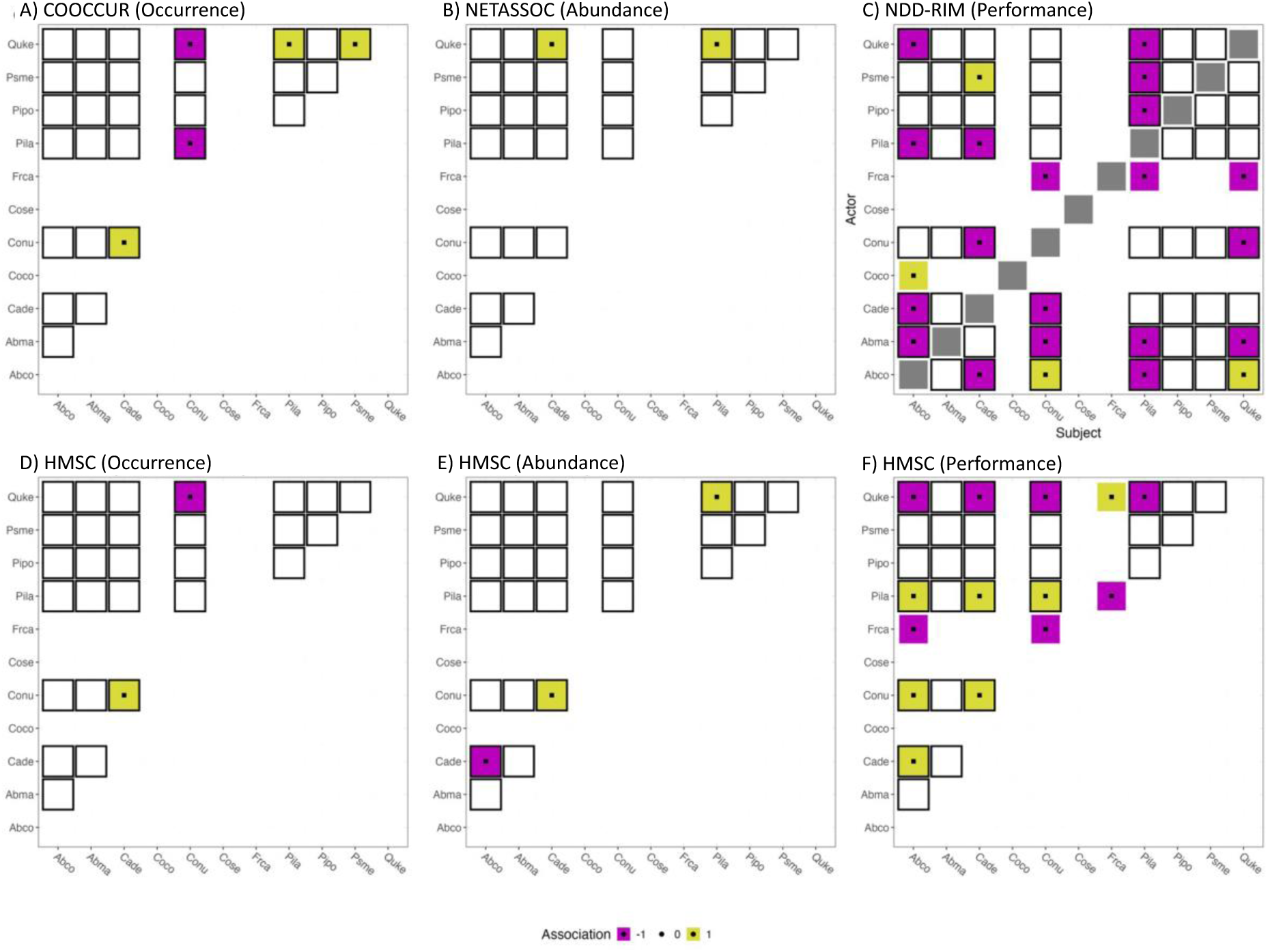
Model Specification Drives Network Inference. Ecological networks inferred at plot scale using the four methods of network inference with differing specifications of biodiversity products (i.e., occurrence, abundance, performance). COOCCUR (A), NETASSOC (B), NDD-RIM (C) represent use-cases for occurrence, abundance, and performance data. HMSC (D-F) can incorporate occurrence (D), abundance (E), or performance (F) data. Inferred associations/interactions have been set to +1 (facilitative), -1 (competitive) or 0 (no link inferred) to ensure comparability. Black outlines highlight species pairings shared across all scales of assessment. Black squares in the network matrices signify statistically significant inferred connections between species. See table S1 for an overview of four-letter abbreviations and corresponding species identities. Note that NDD-RIM (C) infers interactions rather than associations hence rendering non-symmetrical network matrices.

**Figure 3.**
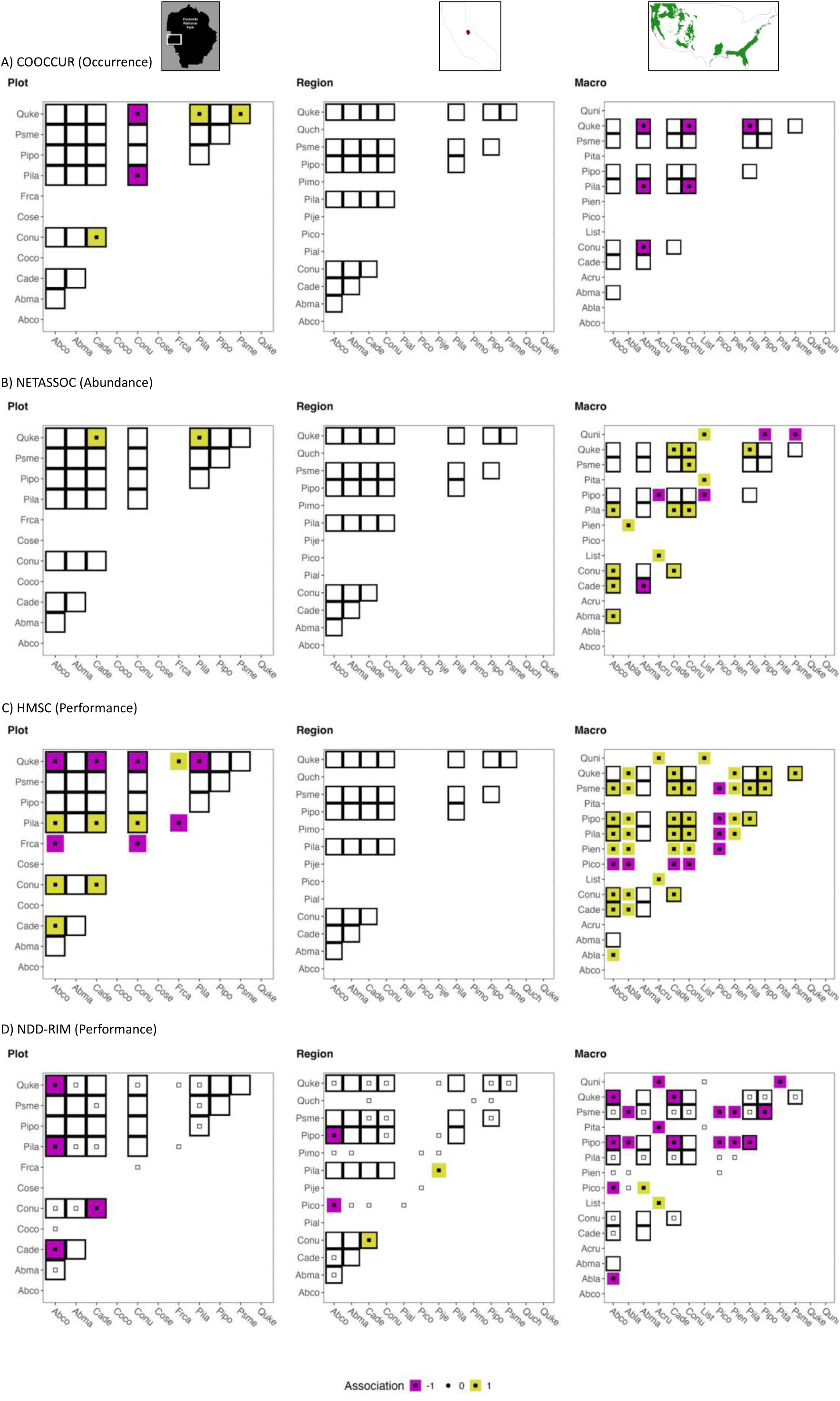
Network Inference is Not Consistent across Approaches or Scales. Contrasting the ecological networks inferred using four separate methods (A-D) reveals a lack of scale-consistency when assessing species-species links across geographic scales (i.e., plot scale – left column, region scale – centre column, and macro scale – right column). Inferred associations have been set to +1 (facilitative), -1 (competitive) or 0 (no link inferred) to ensure comparability. Black outlines highlight species pairings shared across all scales of assessment. Filled dots in the network matrices highlight statistically significant inferred connections between species. Empty dots indicate in NDD-RIM visualisation (D) indicate that only one of the underlying directed links was identified statistically significantly. See table S1 for an overview of four-letter abbreviations and corresponding species identities.

## 3. Results

To assess comparability and consistency of inferred ecological networks across the performance-environment ordination, we focus on plot scale YFDP data (Figure 2) because most data types are available at this scale (Table 1). The corresponding trends observed at plot- scale are mirrored at regional and macro-scales (Figure S4 & Figure S5).

Pairwise inferred links showed consistent patterns across methods for the YFDP dataset (Figure 2). This pattern holds even for different specifications of the HMSC approach (Figure 2D to F). Inference methods focused on performance data (NDD-RIM and HMSC in Figure 2F) identify more statistically significant links between species pairs than those focused only on occurrence or abundance data (Figure 2A, B, D, E) despite notable similarities between occurrence- and abundance-informed HMSC inference at the plot scale.

Inferred network matrices for the eight-species pairings shared across all scales (black outlined tiles in Figure 2 and 3) show no scale consistency even when using the same inference approach. The sign, magnitude, and statistical significance of links change across scales. In the case of HMSC (Figure 3C), we infer a heterogenous network matrix at the plot scale (containing negative and positive statistically significant links) which changes to an exclusively positive, statistically significant link matrix at the macro scale. This scale- inconsistency can also be seen for COOCCUR (Figure 3A) and NETASSOC (Figure 3B), where we fail to detect any statistically significant associations at the regional scale. Lastly, using NDD-RIM, there is little consistency in the sign and magnitude of statistically significantly identified directed links at the plot scale (Figure 2C), regional scale (Figure S4C) and macro scale (Figure S5C). These inconsistencies remain when collapsing opposing directed links inferred by NDD-RIM into undirected links (Figure 3D).

Quantifying inference dissimilarity via decomposition of network-dissimilarities into node-set and link-pattern changes reflects the overall lack of inference consistency across scales and approaches. Dissimilarities of present (positive or negative in sign and non-zero in magnitude) links within scales are high and cannot be traced back to species turnover (Figure 4A) as each network of the same scale consists of the same node set. The same is true for link sets including absent (i.e., non-statistically significant inferred links) as well as present links but to a lesser degree due to sparse inference of present associations (Figure 4B). Instead, these must be attributed to each inference approach’s input data or statistical philosophy. In line with our expected differences due to the use of occurrence, abundance, and performance data, the lowest dissimilarity scores of present links within-scales can be found between HMSC and NDD-RIM outputs. This is likely due to similar information being captured in abundance and performance records (Figure S7). As for abiotic factors governing ecological interactions and their inference, we find comparatively low dissimilarity of present links between HMSC and NETASSOC at the macro scale, for which we hypothesised climatological gradients to play crucial roles. Considering also absent links no such insights can be found. Instead, we identify HMSC as inferring vastly different link motifs as compared to the other approaches. This may be an effect of HMSC’s ingestion of additional data (i.e., functional trait expressions and phylogenetic relatedness).

**Figure 4.**
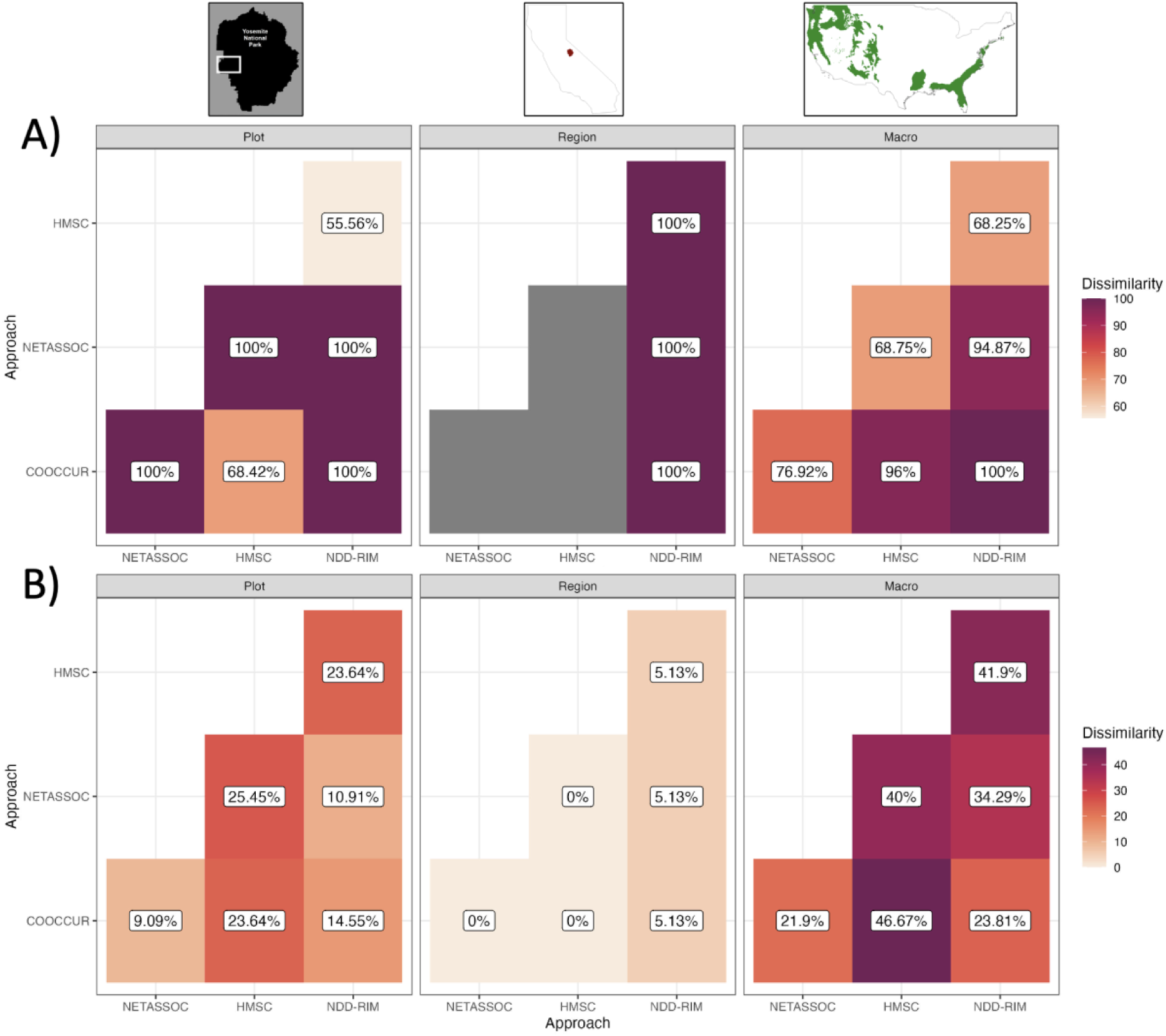
Network Dissimilarity Across Approaches Within Scales Of Assessment. There exists considerably dissimilarity between ecological networks made up of A) statistically significant links and B) statistically significant links and links inferred as absent by the target approaches within each scale of assessment. Notably, inference at region scale (centre panel) was held back by low data availability.

When investigating cross-scale consistencies, we find that dissimilarity of only present links as well as present and absent links is driven by changes in links between species present across the local, regional, and macroecological scale instead of added or removed link structures introduced by species turnover. This dissimilarity may be reduced by large amounts of data (Figure S6). This finding is consistent for all inference methods assessed here with some approaches rendering significantly more similar inferences across scales of assessment than others (Figure S6). Finally, these inconsistencies extend beyond comparisons of individual link structures and lead to a lack of consistency in network topology as described by modularity and node centrality (see supplementary material and Figures S8 & S9).

## 4. Discussion

We have identified a lack of consensus among networks inferred for forest ecosystems using COOCCUR, NETASSOC, HMSC, and NDD-RIM. This discrepancy could be attributed to (1) differences in philosophy of methods (e.g., performance-environment ordination placement or targeted link-types), (2) scale-dependant applicability of different methods and data availability/richness, (3) realised biological interactions change across scales due to differences in species pools, and (4) changes in the ecological processes driving community assembly across scales.

### 4.1. Scale-Consistency & Biological Processes

Biological processes revealed by ecological networks are conditional on the choice and scale of the inference method. The approaches of ecological network inference assessed here focus on representing networks emerging from different ecological processes. While COOCCUR and NETASSOC aim to identify species associations and their outcome on space-utilisation, HMSC is primarily aimed at disentangling community assembly processes with link-inference as a side-product of residuals. In contrast, NDD-RIM considers asymmetries in species associations and aims to identify the effect of one association partner on the performance of the other. In addition, the response-impact model component of NDD-RIM may prove helpful in predicting novel interactions as communities reshuffle, thus giving a valuable set of testable hypotheses for experimental macroecology (Alexander et al. 2016) and management.

Figure 3 and Figure S6 illustrate that 1-1 associations are inconsistent across geographic scales when inferred using the same approach. These scale-inconsistencies are not necessarily a flaw in the methods of network inference but instead a feature of (1) the inference method applicability at varying scales, (2) the scale-dependence of species-interactions, and (3) which biological processes are being captured at specific geographic scales.

Inferred species links can be considered data-driven hypotheses about realised biological interactions (Ovaskainen et al. 2016), which have been demonstrated to change in strength and sign when additional species are introduced/removed (Blanchet et al. 2020; Bartomeus et al. 2021). Our analyses of network link beta diversity suggest that scale-inconsistencies of 1-1 associations are partially driven by variation in realised interactions but cannot be explained by changes in species pools alone. Therefore, while there is a precedent for using ecological networks measured/inferred at one scale to draw conclusions at other scales (Guimarães 2020), our findings indicate that neither 1-1 associations nor whole-network topology measures should be transferred between scales and management decision based on such procedures may prove to be erroneous and detrimental in forest management. For example, we show that ecological networks inferred for individual forest stands such as the YFDP plots would lead to vastly different forest management decisions of temperate conifer forests than decisions based on inferences at whole-biome level. We recommend ecological networks be inferred and interpreted at the scales at which they are to be used for further analyses and decision-making. Additionally, we caution against overconfidence in inferred ecological networks as even within distinct scales can networks vary vastly across approaches and under different parameterisations of the same approach (HMSC inferences in Figures 2, S4, and S5).

### 4.2. Choice of Network Inference Approach Across Scales

Since quantifications of inference performance for methods of ecological network inference under relevant study settings are absent and will likely remain so, we propose to orient choice, use, and interpretation of ecological network inference for forest ecosystems within a framework accounting for the sources of inference discrepancies we have identified. Choice of network inference method should be driven by a combination of criteria. Most importantly, (1) the type of ecological association/interaction which is targeted should be the first selection criterion to be examined. Additionally, our results highlight two additional method selection criteria: (2) biodiversity inputs available to analyses and (3) data integration and availability which we find to be of particular relevance as the scale of study increases. We present these selection criteria and their relation to the assessed network inference approaches in Figure 5.

**Figure 5.**
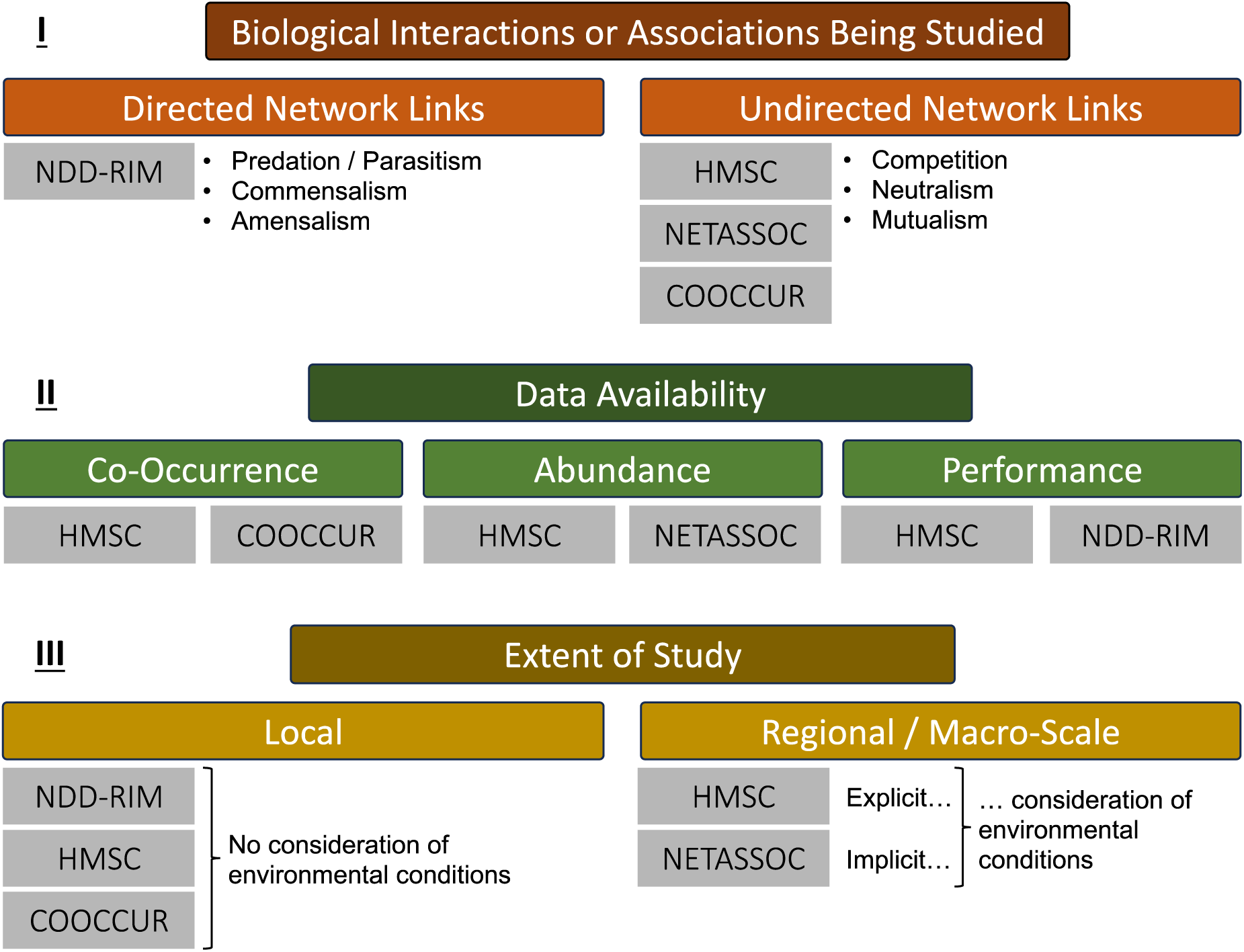
Criteria for Choice of Network Inference Method. Choosing an appropriate approach to study network inference is a complex task that requires careful consideration of three fundamental criteria: I) Which biological connection (association or interaction) is the main interest of the study? Whether inferred networks ought to be directed or undirected should help define the choice of method. II) What data is available? When performance or abundance data is available, it is likely more prudent to use a method of network inference designed to work with this data. III) What is the scale of interest? At regional and macroecological scales, methods of network inference which incorporate environmental information are likely to be the most accurate. Any method of network inference which satisfies the above three criteria for a specific study setting is likely to be more applicable for the desired purpose than a method which does not.

Firstly, ecological network inference approaches have typically not been designed to disentangle asymmetric links. Such directed links characterise biological interactions where each species is influenced differently by the link (e.g., commensalism, predation). In truth, most network inference approaches assume ecological networks are undirected thus suggesting that an ecological network gather either beneficial (e.g., mutualism) or harmful (e.g., reciprocal competition) associations. Due to the ubiquity of these approaches, we have focussed here almost exclusively on such association-based approaches. However, novel approaches such as NDD-RIM have been shown to move past this constraint and infer directed networks, opening the possibilities to study a broader range of association among species (Bimler et al. 2023). Such approaches should be able to account for the entire spectrum of biological associations and interactions (see Figure 5). Thus, while association-based methods may be perfectly suitable to address questions regarding the spatial distribution and co-occurrences of species, performance-based methods are more appropriate for studying the consequences of species interactions/associations on demographic processes. While this capability of inference of directed links lends itself to a wider range of ecological applications, inference of directed links also require more detailed data.

Secondly, while directed-link inference methodology requires abundance/fitness data, inference of undirected links can be carried out using species co-occurrence patterns. This traditional reliance on occurrence data for network inference has been criticised as being too simplistic to measure ecological process outcomes (Blanchet et al. 2020). As seen in Figure 2 and Figure 3, methods of network inference that leverage abundance or performance data identify more statistically significant associations than their occurrence-driven counterparts and thus create more actionable knowledge for application in forest management. We postulate that differences in identified statistically significant ecological associations is partially driven by the amount of biologically relevant information in abundance and performance data compared to occurrence datasets (i.e., quantification of species performances implicitly and explicitly vs. qualitative information). Indeed, performance-based present-links reported the lowest dissimilarity scores (Figure 4). Such differences in information content have already resulted in null model analyses to discriminate community assembly rules being augmented with abundance data (Ulrich and Gotelli 2010). In addition, a recent simulation-based network inference validation framework suggests that the increasing information content along the occurrence-performance spectrum translates directly into higher accuracy of inferred ecological networks (Kusch and Vinton 2023). We suggest that network inference approaches should be subjected to this paradigm shift, thus leveraging performance or abundance data wherever available despite the considerable data collection efforts this entails, particularly at broad geographic scales.

Thirdly, macroecological research has long recognised the relevance of environmental conditions on species presence (Wilkinson et al. 2019), abundance (Johnson and Sinclair 2017), and performance (Coutts et al. 2016). Thus, the scale applicability of ecological network inference depends on whether or not climatological gradients ought to be considered in a given study. Of the four evaluated methods, COOCCUR and NDD-RIM do not account for climatological gradients, while NETASSOC does so implicitly through range expectations, and HMSC does so explicitly through direct use of climate data to quantify contributions of bioclimatic niches and environmental conditions to species performance. By accounting for environmental gradients, which alter species co-occurrence potential, NETASSOC and HMSC address one recurring criticisms of ecological network inference (Blanchet et al. 2020) and may drastically improve network inference accuracy even when leveraging occurrence data (Kusch and Vinton 2023). The effect of climatological data integration into ecological network inference manifests itself in lower network dissimilarity scores for statistically significant inferred links with NETASSOC and HMSC at macroecological scales. The ability to account for climatic data is thus of particular importance when inferring networks at scales for which species’ performances and distributions vary meaningfully as a result of environmental conditions. Note that this scale may be much finer than the macroecological scale depending on the organisms involved.

In addition to including climatological patterns, there are calls to account for the shared evolutionary history of species pairs as well as their functional trait expressions (Ovaskainen et al. 2017; Blanchet et al. 2020). This information is likely to be of particular importance to infer interactions which are evolutionarily conserved (e.g. parasitism) or which can be reliably matched to easily-measurable functional traits (e.g. predation and body size traits). Meta-analyses of networks across geographic scales, regions, link types, and trophic levels have revealed that a combination of these information criteria is required to uncover mechanisms explaining network structure (Xing and Fayle 2021). However, for our study this information was unavailable at some geographic scales (Table 1). Our results clearly show that data sparsity can also affect network inference at the regional scale (Figure 3 and Figure 4), where NDD- RIM was the only capable method. This capability is likely due to the response-impact model component of the method, designed to infer rare/unobserved links. While complex methods like HMSC are seemingly the only ones fit for inferring ecological networks at macroecological scales (although demonstrating markedly dissimilar inference of present and absent links within scales compared to other approaches, Figure 4B) with respect to integration of additional confounding variables, this scale is also where selective data sparsity problem are the strongest, and inference may still be limited as the sampling of relevant information is costly at such scales.

Finally, to illustrate these criteria or our selection framework, consider two study settings necessitating inference of ecological networks: (1) a study of plant-plant relationships within a singular forest stand to evaluate how exotic plants have integrated the native community and (2) a study of plant-frugivory networks at several locations around the Globe to predict range shifts in response to global warming. For study (1), it is of pivotal relevance to select a method of network inference which is capable of rendering directed links as competition between plants is rarely symmetrical (e.g., NDD-RIM, for example). Since the study system is small in extent, climatological gradients will likely play a minor role in altering occurrence, abundance or performance of forest-stand organisms over their biological interactions. Thus, choosing an approach without consideration of environmental conditions is likely adequate (e.g., COOCCUR, NDD-RIM, or HMSC and NETASSOC without environmental inputs). Finally, if abundance or performance data is available or can be sampled, it would be prudent to select an approach capable of using such data (e.g., NDD-RIM, NETASSOC or HMSC). Ultimately then, for this study purpose, one should select (from the pool of approaches demonstrated here) NDD-RIM for network inference as it satisfies all selection criteria and, most importantly, infers directed links which are required to express interaction effects from natives on exotic plants and vice versa. As for study (2), we are dealing with a broad scale of study at which consideration of climatological gradients will likely be highly relevant for accurate network inference and answering our research questions. Therefore, we may elect to use either HMSC or NETASSOC for inference in this setting. This aligns with the requirement of needing to infer mutualistic interactions which may be expressed as undirected links (although a case could be made that directed links would contain more information even in such a setting) which would also predispose a choice of HMSC or NETASSOC. The final choice between which of these two approaches to use will likely be informed by whether phylogenetic relatedness of species or their functional trait expressions are known which would subsequently give an advantage to HMSC because it can readily use such data. Ultimately, when arriving at an impasse in choosing between competing methods of network inference, one may wish to employ novel simulation-based network inference validation frameworks with study specific parameterisation to quantify network inference performance a priori (Kusch and Vinton 2023). Nevertheless, it is unlikely that any single network inference approach will render a fully accurate counterpart to real-world ecological interactions/associations. Instead, to maximise inference accuracy at the cost of a deeper understanding of factors structuring ecological networks, it may be prudent to combine different inference approaches in an ensemble framework to establish more robust network inference while also accounting for the different ecological processes which disparate inference approaches are centred on.

## 5. Conclusion

Methods of network inference provide a valuable tool to gain a deeper understanding of ecological systems. They can suggest potential biological interactions and associations that may be worth exploring more completely depth and that would not have been considered otherwise (e.g., because of the size of communities that can include many potential and realised interactions). We show however, that scale inconsistencies are considerable for pairwise species links. In the absence of careful simulation-based assessment of network inference performance for specific study criteria, we argue that the choice of method needs to consider: (1) biological mechanisms which are identifiable using network inference at the scale of study, (2) alignment of link-interpretation with research questions, (3) data availability and information content at the geographic scale of study, and (4) applicability of the network inference method at the geographic scale of study.

## Acknowledgements

This research was supported by the Aarhus University Research Foundation Start-up Grant (grant no. AUFF-2018-7-8) to AO. AO considers this a contribution to the Center for Ecological Dynamics in a Novel Biosphere (ECONOVO), funded by Danish National Research Foundation (grant DNRF173). Additionally, this work was supported by the European Union’s Horizon Europe research and innovation programme under grant agreement No. 101057437 (BioDT, https://doi.org/10.3030/101057437).

## Author contributions

EK and AO conceptualised this study. EK curated data and led the analysis. EK created all R scripts necessary for the analyses. NDD-RIM integration was aided by MB. Results were interpreted by EK and AO. YFDP data was provided by JAL. FGB provided critical feedback and aided writing efforts. Manuscript writing was led by EK. All authors contributed critically to the drafts and gave final approval for publication.

## Data Availability

This submission uses novel code, provided at https://github.com/ErikKusch/Ecological-Network-Inference-Across-Scales. Download queries for raw FIA and ERA5-Land data are specified in the R scripts (FIA data at https://apps.fs.usda.gov/fia/datamart/datamart.html; ERA5-Land data at https://cds.climate.copernicus.eu/cdsapp#!/dataset/reanalysis-era5-land-monthly-means). Raw YFDP data was downloaded from the Smithsonian ForestGEO data portal using this form http://ctfs.si.edu/datarequest/index.php/request/form/6 and specifying the following query: “full data predating fire-event in 2013”. All subsequent handling of raw data is detailed in the R scripts.

