## Supplement for "Inconsistencies of empirical ecological network inference are governed by considerations of statistical approaches and dimensions of input data"

TITLE:

Erik Kusch[1,2,3] (0000-0002-4984-7646)

Malyon D. Bimler[4] (0000-0003-0059-2360)

F. Guillaume Blanchet[5,6,7] (0000-0001-5149-2488)

James A. Lutz[8] (0000-0002-2560-0710)

Alejandro Ordonez[2,3,9] (0000-0003-2873-4551)

CORRESPONDING AUTHOR:

Erik Kusch;

### **THE OCCURRENCE-PERFORMANCE SPECTRUM**

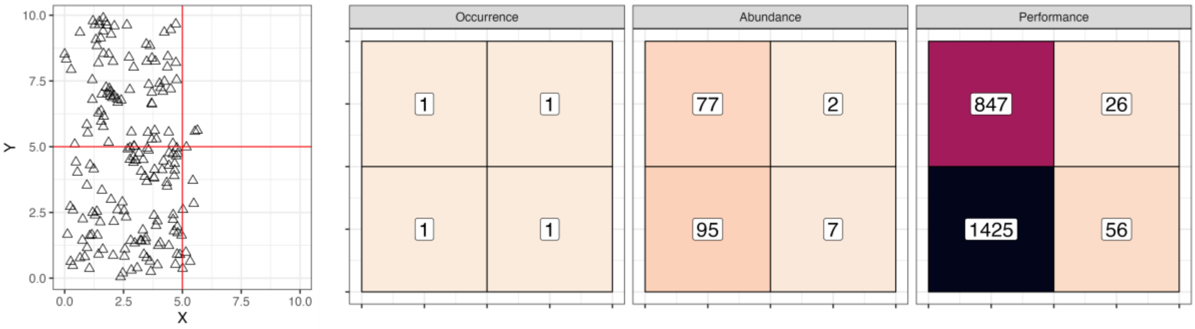


**FIGURE S1 – THE CO-OCCURRENCE-PERFORMANCE SPECTRUM OF NETWORK INFERENCE INPUTS** – Left panel: spatial arrangement of individuals belonging to one species (lines indicate site borders). Right panel: species-by-site representations of biodiversity data corresponding to individuals in left panel. Note that this is a conceptual visualisation and not sourced from the empirical data used in this study.

### NETWORK INFERENCE APPROACHES

***Probabilistic Species Co-Occurrence Analysis in R (COOCCUR)***: This probabilistic approach identifies species associations from co-occurrence patterns by analytically estimating the probability of each species-pair co-occurring either less or more often than random, assuming species occurrence probability at each site equals the observed frequency among all sites (Veech 2013). The calculated probabilities of co-occurrence of each species-pair being greater/smaller than that random are subsequently used as p-values to identify the statistical significance of each species-species association.

***Inference of Species Associations from Co-Occurrence Data (NETASSOC)***: NETASSOC compares an observed species-by-site matrix (either of abundances or occurrence records) against null model-derived species-by-site matrices to identify non-random species association patterns (Morueta-Holme et al. 2016). In contrast to COOCCUR, these null models are not strictly limited to random assemblages. Instead, using species range estimates for each species contained within the observed species-by-site matrix, NETASSOC can establish a null expectation for species occurrence at sites incorporating bioclimatic gradients but excluding biotic associations that are subsequently estimated through NETASSOC. The statistical significance of identified species-species associations is determined by two-tailed p-values for each species pair and corrected for multiple comparisons.

***Hierarchical Modelling of Species Communities (HMSC)***: This approach is a flexible hierarchical Bayesian joint species distribution model that identifies community assembly processes (Ovaskainen et al. 2017), which underpin both potential and realised ecological associations (Montesinos-Navarro et al. 2018). HMSC combines occurrence, abundance, or performance species-by-site matrices with environmental information at each site to identify environmental community assembly filters. The species niches identified by HMSC can be further informed by each species' phylogenetic relatedness and functional trait expression. From this information, HMSC fits a species distribution model based on observed species-by-site data. It then treats residual species-species correlations as biotic filtering processes like species-species associations. Statistical significance of identified species associations is then assessed using credible intervals of their posterior distributions (Opedal and Hegland 2020).

***Neighbour-Density Dependent - Response Impact Model (NDD-RIM)***: This approach differs from other methods in its' aim to quantify species-specific effects on growth/survival rather than co-occurrence probabilities (Bimler et al. 2022). Thus, links between nodes identify focal species' performance changes due to interaction partners' abundance. These directed links between the network nodes are quantified by leveraging performance at sites. Links between nodes are modelled through a Neighbour-density dependent (NDD) model component in which performance metrics are regressed against the identity and abundance of neighbouring species. Latent variables capturing the response (whether a species responds, on average, positively or negatively to neighbouring species) and effect (the average change in all focal species' performance exerted by a given species) parameters of each focal species are used to infer rare/unobserved interactions. We refer to this model component as Response-Impact-Model (RIM). Since this method is built on a Bayesian framework, posterior distribution intervals can be used for assessing the statistical significance of the identified species interactions.

NDD-RIM has been developed for individual-level data which is not available at our larger scales of assessment (i.e., FIA data). Therefore, we used a modified version of NDD-RIM at these scales to account for potential correlations between observations of performance metrics obtained for the same plot. The modified version included a random effect for plot, and we employed the 'sum-to-zero' approach detailed in Ogle & Barber, 2020 to avoid identifiability issues.

### DATA & ANALYSIS WORKFLOW

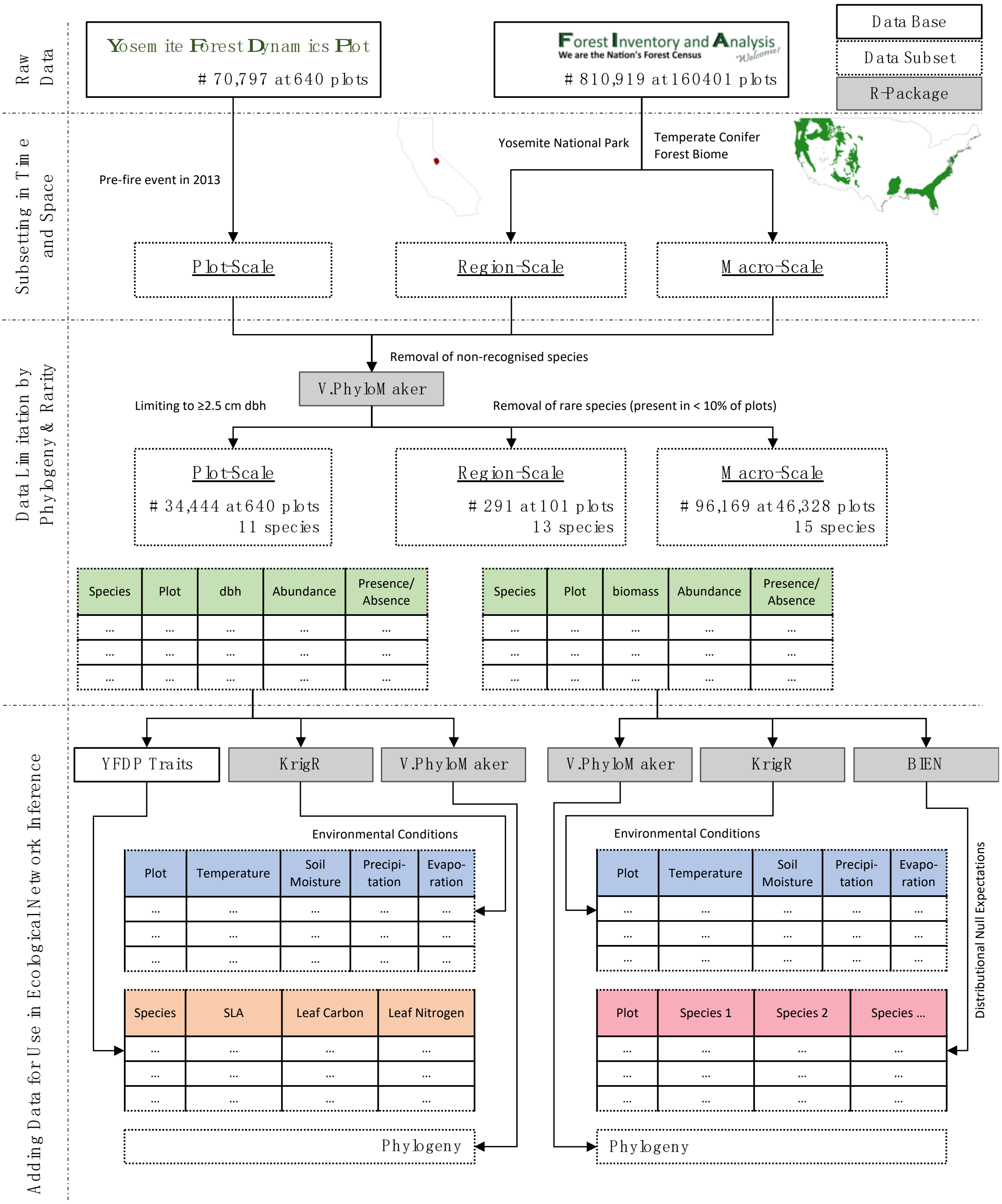


**FIGURE S2 – DATA HANDLING WORKFLOW** – Data for this analysis was obtained from a variety of data sources and subsequently divided into observations at plot, regional, and macro scale.

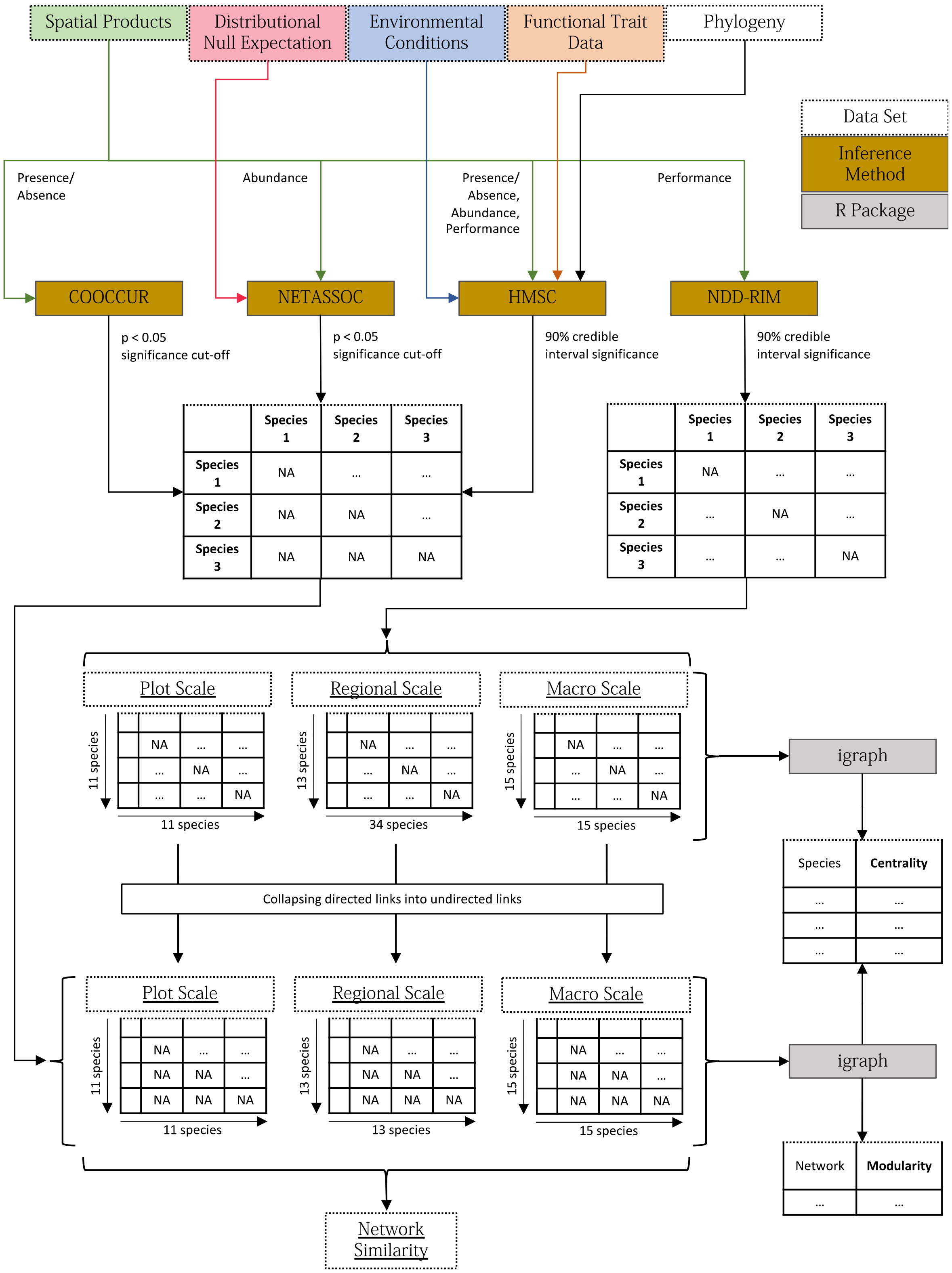


**FIGURE S3 – NETWORK INFERENCE WORKFLOW AND CALCULATION OF NETWORK TOPOLOGY METRICS** – An overview of data source usage by network inference methods, output (i.e., association or interaction matrix). Note that interaction matrices obtained from NDD-RIM are used for centrality calculation but collapsed into association matrices for calculations of modularity.

### SPECIES IDENTITIES

**TABLE S1 – SPECIES IDENTITIES** – Network visualisations use a four-letter code to denote species identities. These four-letter codes correspond to binomial nomenclature as shown here.

| **Abbreviation** | **Species Name** |
|---|---|
| Abco | *Abies concolor* |
| Abma | *Abies magnifica* |
| Cade | *Calocedrus decurrens* |
| Conu | *Cornus nuttallii* |
| Cose | *Cornus sericea* |
| Coco | *Corylus cornuta* |
| Frca | *Frangula californica* |
| Pila | *Pinus lambertiana* |
| Pipo | *Pinus ponderosa* |
| Psme | *Pseudotsuga menziesii* |
| Quke | *Quercus kelloggii* |
| Pial | *Pinus albicaulis* |
| Pico | *Pinus contorta* |
| Pije | *Pinus jeffreyi* |
| Pimo | *Pinus monticola* |
| Quch | *Quercus chrysolepis* |
| Abla | *Abies lasiocarpa* |
| Acru | *Acer rubrum* |
| List | *Liquidambar styraciflua* |
| Pien | *Picea engelmannii* |
| Pita | *Pinus taeda* |
| Quni | *Quercus nigra* |

### Model Specification Drives Network Inference

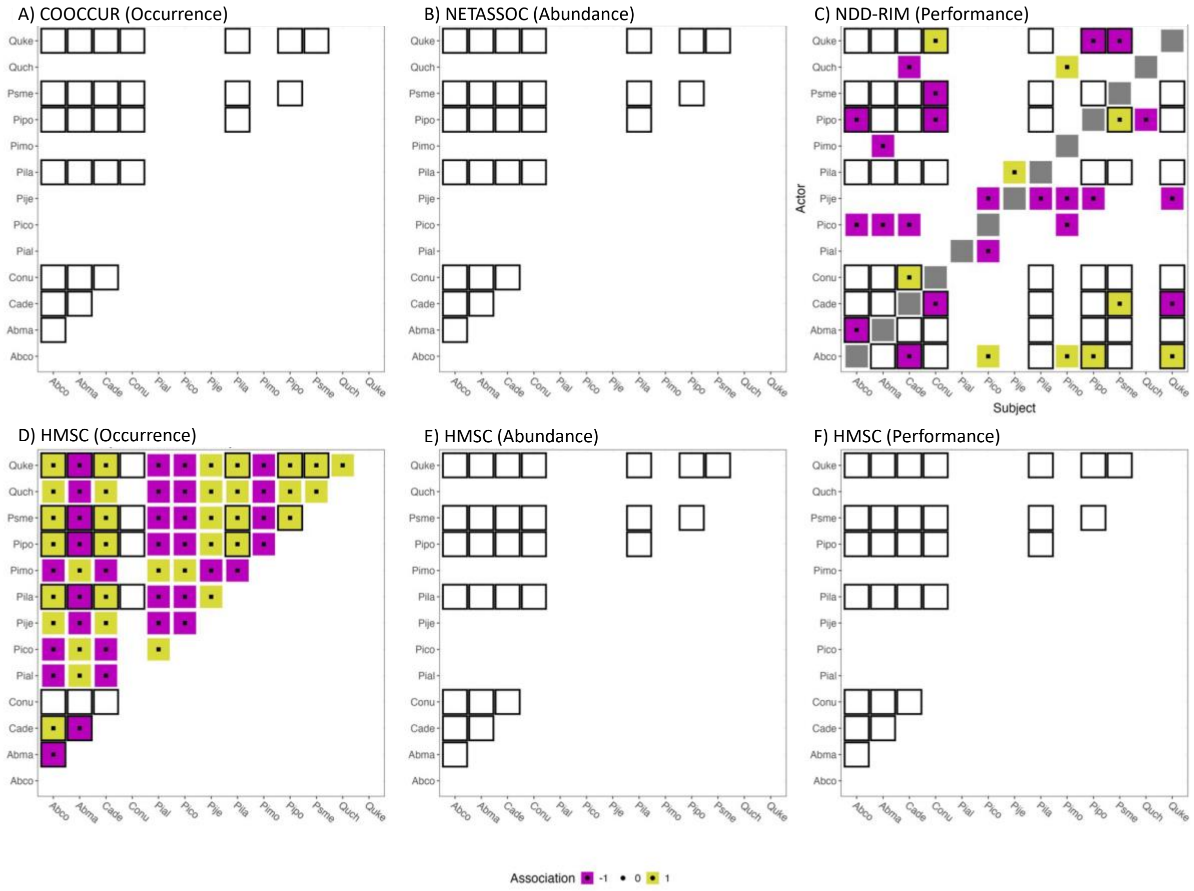


**Figure S4 - Model Specification & Network Inference at Regional Scale** – Ecological network adjacency matrices inferred at regional scale using the four methods of network inference with differing specifications of spatial products. Inferred associations/interactions have been set to +1 (facilitative), -1 (competitive) or 0 (no link inferred) to ensure comparability. Black squares in the network matrices signify statistically significant inferred connections between species. See table S1 for an overview of four-letter abbreviations and corresponding species identities.

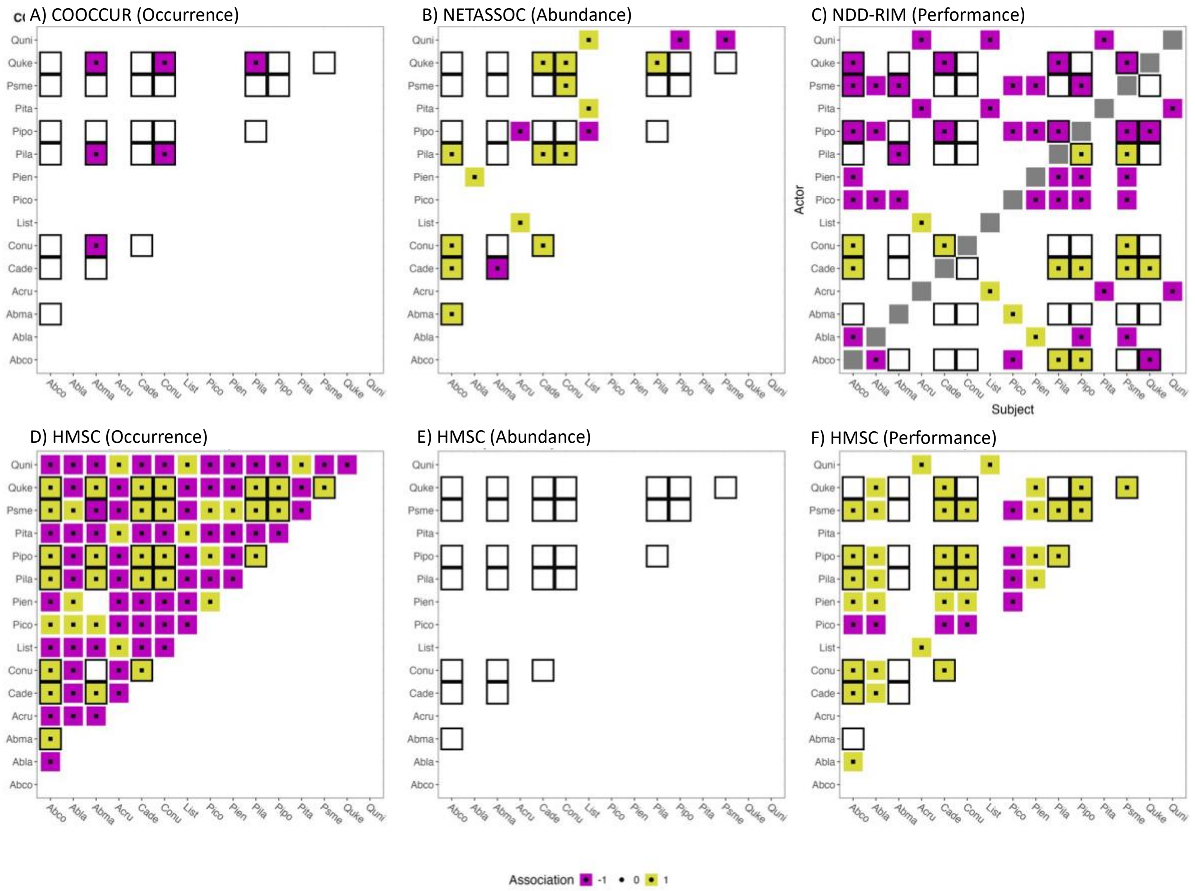


**Figure S5 - Model Specification & Network Inference at Macro Scale** – Ecological network adjacency matrices inferred at macro scale using the four methods of network inference with differing specifications of spatial products. Inferred associations/interactions have been set to +1 (facilitative), -1 (competitive) or 0 (no link inferred) to ensure comparability. Black squares in the network matrices signify statistically significant inferred connections between species. See table S1 for an overview of four-letter abbreviations and corresponding species identities.

### DISSIMILARITY OF INFERRED NETWORKS

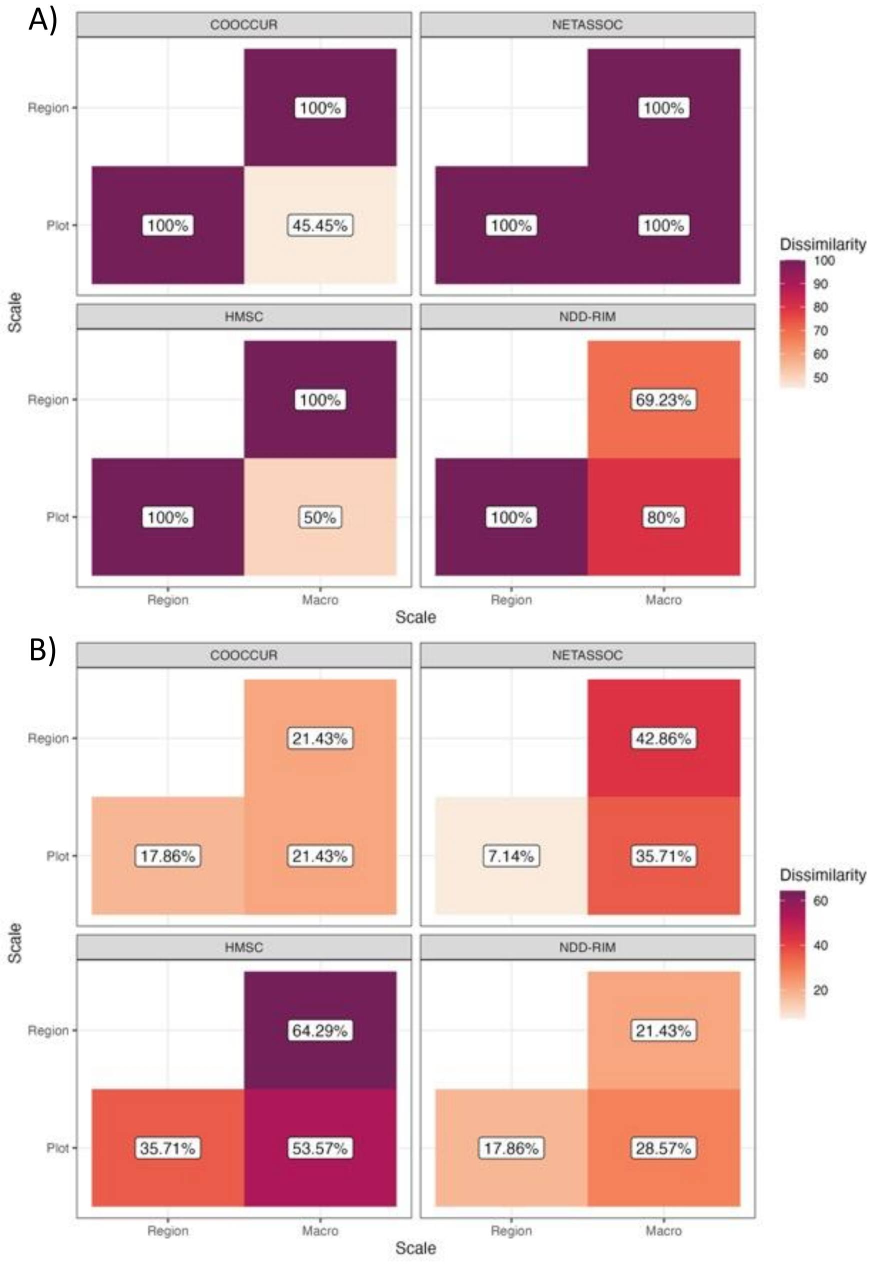


**FIGURE S6 - NETWORK DISSIMILARITY ACROSS SCALES OF ASSESSMENT WITHIN APPROACHES** – Network dissimilarity scores are high when contrasting networks inferred by the same network inference approach across the three distinct scales of assessment of our study. A) Comparing only statistically significant links, dissimilarity scores between plot and macro-scale are lower than those between regional-macro and regional-plot scales which is probably due to the low amount of data for use in at the regional scale in our study. B) Comparing statistically significant links and links inferred as absent, we find large cross-scale inconsistencies for HMSC inference and NETASSOC but reduced cross-scale inconsistencies for COOCCUR and NDD-RIM.

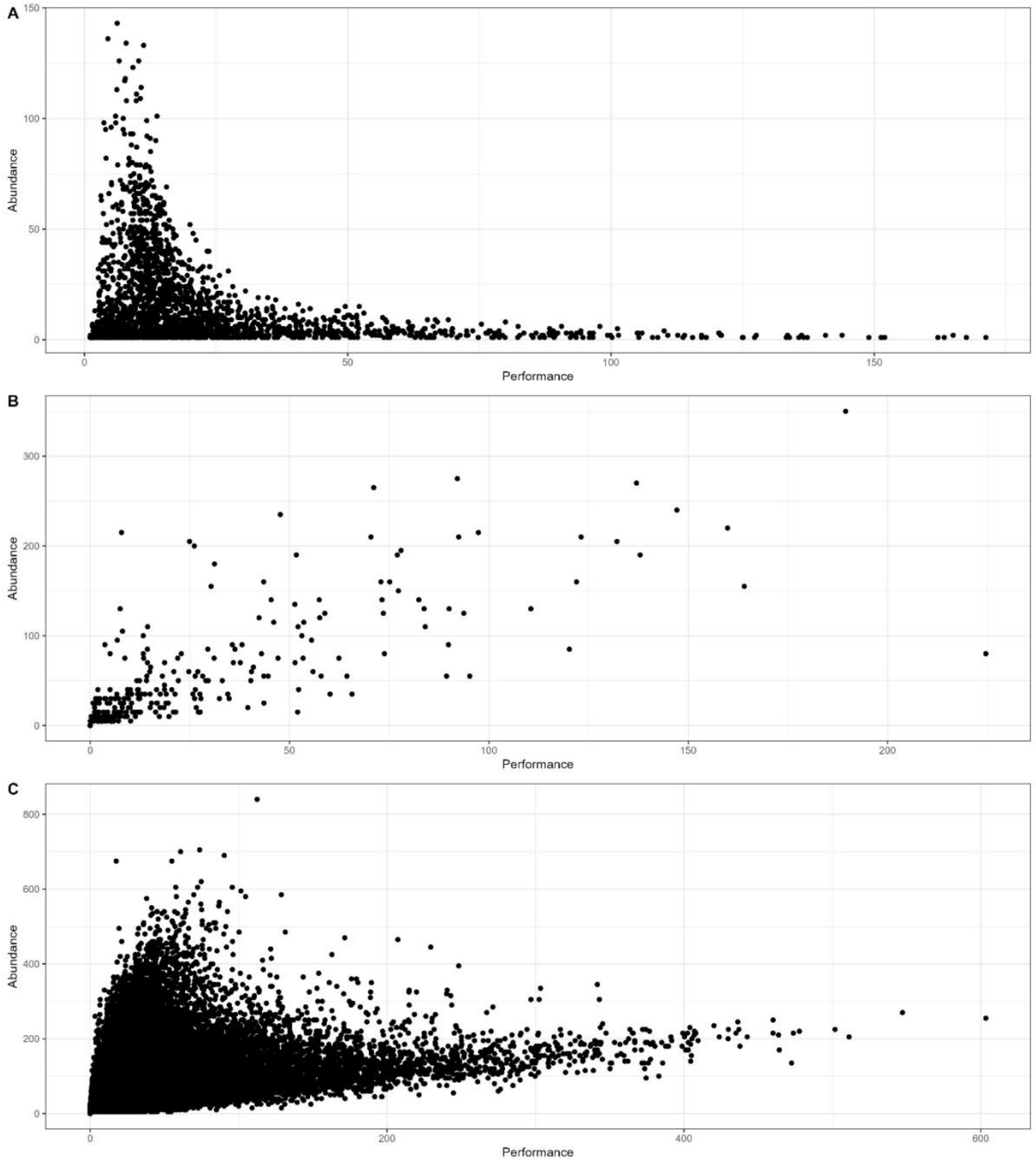


**Figure S8 – Similarity of Abundance and Performance Data across scales of assessment** – Abundance (y-axis) and Performance (x-axis) data used at the (A) local, (B) regional and (C) macro scale in this study. As scale increases, abundance and performance (as measured through species-specific biomass) become more aligned and one may be used as a proxy of the other for the purpose of pairwise species-association inference.

### NETWORK TOPOLOGY METRICS

We used three network topological attributes (modularity, nestedness, and node centrality; Table S3) linked to ecosystem stability to contrast inferred networks yielded by the four implemented methodologies. Just like the computation of network similarity, this analysis also used only those links which were identified statistically significantly. Topological attributes of inferred association network were estimated using the igraph (Csardi and Nepusz 2006) R package. To ensure comparability of methods, we collapsed opposing links in NDD-RIM networks into mean links when computing topological metrics.

**TABLE S2 - NETWORK TOPOLOGY METRICS** – Network topology metrics can be classified as either describing individual network nodes (node-level metrics) or the entire network (network-level metrics) (Bascompte and Jordano 2007, Morueta-Holme et al. 2016). The list of topology metrics presented here is aimed at giving an overview of often-used metrics, their topology-level, and definition. As such, this listing is non-exhaustive.

| Metric | Level | Definition | Reference |
|---|---|---|---|
| **Centrality** | Node | Sum of absolute link weights/magnitudes connected to individual nodes (also referred to as “weighted degree” or “node strength”). | (Farine and Carter 2022) |
| **Degree** | | Number of realised links to other nodes | (Bascompte and Jordano 2007, Morueta-Holme et al. 2016) |
| **Modularity** | Network | Level of compartmentalisation of network into well-connected assemblages and nodes which are sparsely connected with other hubs within the same network | (Thébault and Fontaine 2010, Morueta-Holme et al. 2016, Landi et al. 2018) |
| **Nestedness** | | Level of sharing of interaction partners between generalist and specialist species | (Thébault and Fontaine 2010, Landi et al. 2018) |

When assessing consistency in ecological network attributes, we focused on the sub-networks containing the eight species shared across all scales. We did so to avoid biases in topological metrics introduced by changes in network size (Poisot et al. 2012). We evaluated differences in network topology metrics both at a whole network and node-level (see Table S3). To facilitate the comparison of centrality scores across networks belonging to different inference methods, we subsequently transformed all centrality scores to a range of 0 to 1 per network.

Inferred network modularity, much like the underlying network matrices (Figure 3 for undirected NDD-RIM networks), shows no consistency across methods or scales (Figure S8). For example, across scales, whereas the HMSC network at plot scale shows relatively high modularity, the corresponding 8-species network at macro scale exhibits much lower modularity. Contrasting across methods at the plot scale shows variable modularity estimates. Specifically, COOCCUR identifies the lowest modularity, NETASSOC renders a network of no separated modules (purely positive associations, see Figure 2B), HMSC identifies relatively strong modularity, and NDD-RIM identifies the strongest modularity. Note that we calculated modularity based only on statistically significant associations identified by the inference methods.

While there exists little consensus regarding potential keystone species classification across scales, there is general agreement on node importance across evaluated methods. Specifically, three out of four approaches agree on the high ecological importance of *Pinus lambertiana* (Pila) and *Abies concolor* (Abco) in the ecological networks at the macro scale (see Figure S9). Furthermore, at both the local and macro scale, *Abies concolor*, *Pinus lambertiana*, show high centrality scores for HMSC, implying some scale-consistency of this method. The same is true for *Pinus ponderosa* at regional and macro scale for NDD-RIM.

Network topology metrics, which have been linked to ecosystem resilience and functioning (Thébault and Fontaine 2010). Our analyses highlight that whole-network topology metrics are also subject to scale inconsistencies (see Figure S8).

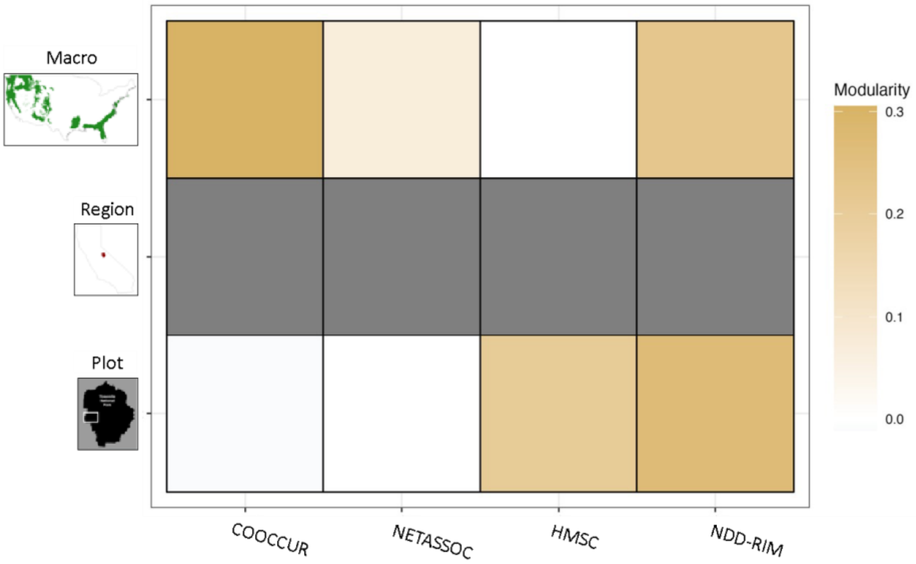


**FIGURE S8 – MODULARITY OF NETWORKS INFERRED ACROSS METHODS AND SCALES IS NOT CONSISTENT** – Inferred ecological network modularity across inference methods (columns) and scales (rows). Note that COOCCUR, NETASSOC and HMSC identified no statistically significant connections at some scales. No modularity values could be calculated for these cases. Additionally, when network modules are perfectly separated (i.e., NDD-RIM at regional scale), modularity cannot be calculated using the spinglass method.

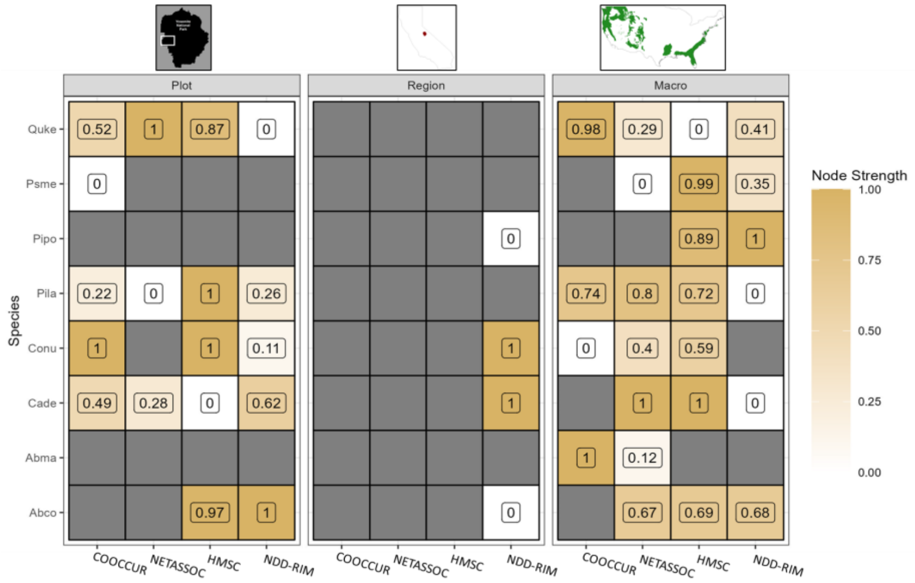


**FIGURE S9 – CENTRALITY OF NODES CONTAINED IN NETWORKS INFERRED ACROSS METHODS AND SCALES** – Node centrality for species shared across scales in our analyses. Centrality was calculated for each inferred ecological network and subsequently standardised to a data range of 0 to 1 to enable comparison of networks across scales and methods. Note that COOCCUR, NETASSOC and HMSC identified no statistically significant connections at some scales resulting in no centrality scores being calculated for these cases. See table S2 for an overview of four-letter abbreviations and corresponding species identities.

While we caution against transferring entire inferred ecological networks or their topologies across spatial scales, our analysis did identify scale consistency for sub-network topology metrics. Specifically, measurements of individual species' importance in shaping their communities (i.e., centrality) remain consistent across geographic scales and, to some extent, even network inference method. For example, *Cornus nutallii* was identified consistently as one of the most central nodes among our three distinct geographic scales. At the macro-scale and focusing on methodology applicable for this scale (i.e., HMSC and NETASSOC), *Calocedrus decurrens* and *Pinus lambertiana* have been identified as the most central nodes in their respective ecological networks. This pattern indicates that some species retain their ecological relevance irrespective of geographic scale or network inference method.
